# Stability of ripple events during task engagement in human hippocampus

**DOI:** 10.1101/2020.10.17.342881

**Authors:** Yvonne Y. Chen, Daniel Yoshor, Sameer A. Sheth, Brett L. Foster

**Affiliations:** Department of Neurosurgery; 1 Baylor Plaza, Houston, Texas, 77030, USA; Department of Neuroscience, Baylor College of Medicine. 1 Baylor Plaza, Houston, Texas, 77030, USA; Department of Neurosurgery, University of Pennsylvania, 3400 Spruce Street, Philadelphia, Pennsylvania, 19104, USA

## Abstract

Periods of cognitive disengagement, such as rest or sleep, are thought to support the progressive consolidation of episodic memories. During these states, the hippocampus displays transient high-frequency oscillatory bursts, known as ripples, which are thought to promote interactions with the neocortex, consolidating memory traces. More recent findings have suggested ripples in the human hippocampus may also occur during task engagement, particularly for tasks requiring episodic memory processes. However, it is unclear if hippocampal ripples occur during other cognitive states or whether ripple properties are modulated by specific types of task demands. In addition, identifying genuine hippocampal ripple events in the human brain can be methodological challenging. To address these questions, we used intracranial recordings from the human hippocampus to quantify ripple events across perceptual, memory and resting task states. Using spectro-temporal identification of hippocampal ripples, we observed highly similar ripple event properties across tasks, with a modest yet significant increase in ripple properties (rate, duration & amplitude) during resting task states. These ripple event attributes did not differ between hemisphere, nor across or within the time of day examined. Supporting data further highlighted that while hippocampal ripples occurred during all task states, these rates were typically lower than that observed during sleep. Together, these findings highlight that hippocampal ripples occur consistently, but sparsely, during a broad range of cognitive task states. Such findings may be incorporated into existing models of systems consolidation, whereby hippocampal ripples help to initially establish latent memory traces.

## Introduction

Long term memory formation is thought to be supported by consolidation processes occurring during ‘offline’ states, the most prominent being sleep (Klinzing et al., 2019). During sleep, a host of electrographic signatures are reliably observed in the hippocampus and neocortex, which are thought to underlie systems memory consolidation (Diekelmann and Born, 2010; Joo and Frank, 2018; Klinzing et al., 2019). Specifically, during non-rapid eye movement (NREM) sleep periods, high-frequency oscillations (~80-120 Hz in humans), known as ripples or sharp-wave ripples, are observed in the hippocampus (Bragin et al., 1999; Buzsaki, 2015; Staresina et al., 2015; Helfrich et al., 2019; Jiang et al., 2019). Furthermore, hippocampal ripples display a temporal coordination with other canonical NREM sleep signatures including spindles (~12-16 Hz) and slow oscillations (<1 Hz) (Sirota et al., 2003; Staresina et al., 2015). As part of this orchestrated dynamic, hippocampal ripple events are thought to support interactions with the neocortex as part of a two-stage model for systems consolidation (Klinzing et al., 2019). Under this view, newly encountered information is initially encoded by the hippocampus, but progressively becomes consolidated in the neocortex (Diekelmann and Born, 2010; Dudai et al., 2015; Chen and Wilson, 2017).

While these consolidation processes are most pronounced during offline states, when cognitive engagement is thought to be reduced, recent findings build on prior work suggesting that similar electrographic signatures, particularly ripples, can be observed during awake states (Axmacher et al., 2008). Most recently, it has been reported that ripple events occur in the human hippocampus during the performance of episodic memory tasks (Norman et al., 2019; Vaz et al., 2019). Interestingly, ripples were observed at rates equal or greater than that typically observed during NREM sleep (Jiang et al., 2020; Ngo et al., 2020). Furthermore, hippocampal ripple events were associated with neocortical reinstatement activities and successful memory retrieval (Norman et al., 2019; Vaz et al., 2019). These findings suggest that hippocampal ripples may occur during active memory behavior, supporting hippocampal-neocortical interactions, similar to that observed during sleep (Joo and Frank, 2018).

In light of these findings, it is important to ascertain the specificity of online ripple events for memory behaviors typically associated with the hippocampus and other medial temporal lobe (MTL) regions. At present, there is limited evidence regarding how the occurrence of hippocampal ripples, and their attributes, differ across cognitive task states, including those not typically associated with episodic memory (Buzsaki, 2015). Ripple attributes, including rate, duration, and frequency, likely modulate the mechanistic impact of these events. Recently, Norman et al. (2019) reported that across different stages of a memory recall paradigm ripples events were similar in amplitude and frequency, but differed slightly in rate, suggesting a general stability of attributes across online task states.

In order to sensitively assess ripple activity in human hippocampus, direct invasive recordings are optimal. However, such recordings only typically occur within the context of monitoring being performed for the neurosurgical treatment of refractory epilepsy. This clinical context produces several confounds for the study of ripple activity that require attention, as hippocampal inter-ictal spikes associated with epileptogenic tissue produce transient amplitude increases over a broad frequency range, including the ripple band (Jiang et al., 2020). Therefore, careful consideration of these artifacts and their spectral signatures is required in order to confidently identify genuine hippocampal ripples and in turn their attributes across cognitive tasks.

In the current study, we examined hippocampal ripple attributes across cognitive tasks in 17 subjects undergoing invasive monitoring for epilepsy surgery. We report a striking stability of ripple attributes (rate, duration, amplitude & frequency) across perceptual and mnemonic tasks, while resting conditions showed a slight elevation in ripple rate, duration and amplitude. In addition, we observed no significant difference of these attributes based on anatomical factors within the hippocampus. Furthermore, we find these attributes were stable throughout the time of day and proximity to electrode implantation. However, we report that hippocampal ripples occur at significantly lower rates than previously reported. Indeed, we observed ripple rates, that while stable across online tasks/states, were significantly lower than those observed during offline resting and NREM sleep (Buzsaki, 2015). One factor likely impacting this difference is the criterion employed for ripple detection during online task performance. Finally, we discuss these results within the broader context of how task related ripple events, while at lower rates, may serve to establish hippocampal-neocortical dynamics that are later elevated during sleep to support memory consolidation.

## Materials and Methods

### Human Subjects

Intracranial recordings from the human hippocampus were obtained from 17 subjects (S1-17, 7 male, mean age 34.7 yrs, range 20 – 59 yrs; see Table 1) undergoing invasive monitor as part of their treatment for refractory epilepsy at Baylor St. Luke’s Medical Center (Houston, Texas, USA). Recordings were performed using stereo-electroencephalography (sEEG) depth electrodes (PMT Corp., MN, USA; Ad-Tech Medical Instrument Corp., WI, USA). All experimental protocols were approved by the Institution Review Board at Baylor College of Medicine (IRB protocol number H-18112), with subjects providing verbal and written consent to participate in this study.

**Table 1.** Subject and electrode information. For each subject (1-17) demographic and experimental information is reported in the following order: Sex (Male/Female); Age at time of experiment (years); Experimental tasks performed: Perception (P), Memory (M), Resting (R), other additional Perception tasks and sleep recording; Identified electrode count within hippocampus and hemisphere (Left/Right); Included electrode count within hippocampus and hemisphere (Left/Right).

| Sub | Sex | Age | Task Performed | Elecs. in Hipp. |  |  | Included elecs. |  |  |
| --- | --- | --- | --- | --- | --- | --- | --- | --- | --- |
|  |  |  |  | Tot. | L | R | Tot. | L | R |
| 1 | F | 37 | P, M | 8 | 5 | 3 | 2 | 0 | 2 |
| 2 | F | 32 | P, M | 3 | 3 | - | 3 | 3 | - |
| 3 | F | 59 | P, M | 9 | 9 | - | 4 | 4 | - |
| 4 | M | 53 | P, M | 10 | 6 | 4 | 3 | 3 | 0 |
| 5 | F | 32 | P, M | 1 | 1 | - | 1 | 1 | - |
| 6 | M | 25 | P, M | 13 | 9 | 4 | 13 | 9 | 4 |
| 7 | M | 26 | P, M | 3 | 3 | - | 1 | 1 | - |
| 8 | F | 33 | P, M | 7 | 4 | 3 | 4 | 4 | 0 |
| 9 | F | 27 | P, M | 14 | 7 | 7 | 10 | 6 | 4 |
| 10 | F | 39 | P, M, R | 12 | 5 | 7 | 10 | 3 | 7 |
| 11 | F | 40 | P, M | 7 | 3 | 4 | 4 | 0 | 4 |
| 12 | F | 33 | P, M, R, additional P tasks, sleep | 6 | - | 6 | 6 | - | 6 |
| 13 | M | 22 | P, M | 5 | 5 | - | 4 | 4 | - |
| 14 | M | 39 | P, M | 4 | 3 | 1 | 4 | 3 | 1 |
| 15 | M | 30 | P, M | 5 | - | 5 | 5 | - | 5 |
| 16 | F | 20 | P, M, R | 12 | 12 | - | 6 | 6 | - |
| 17 | M | 43 | P, M | 8 | 3 | 5 | 2 | 1 | 1 |
|  |  |  |  | **127** | 73 | 49 | **82** | 48 | 34 |

### Electrode localization and selection

To identify electrodes located within the hippocampal formation, a post-operative CT scan was co-registered to a pre-operative T1 anatomical MRI scan for each subject, using FSL and AFNI (Cox, 1996; Dale et al., 1999). The volume location of each electrode was identified by clear hyperintensities on the aligned CT using AFNI and visualized using iELVis software functions in Matlab (v2016a, MathWorks, MA, USA) (Groppe et al., 2017). Within subjects, electrodes located within or at the margin of the hippocampal formation were identified, along with electrodes located within the white matter for each depth electrode targeting the medial temporal lobe. As detailed below, white matter electrodes were employed to re-reference hippocampal sites within each probe. Each participant performed several experimental tasks during sEEG recordings.

### Experimental Design and Statistical Analysis

Data analysis focuses primarily on three tasks all performed at the bedside in a quiet and dimmed patient room (described below). Tasks were presented on an adjustable monitor (1920 x 1080 resolution, 47.5 x 26.7 cm screen size, connected to an iMac running OSX 10.9.4) at a viewing distance of 57 cm. Psychtoolbox functions (v3.0.12) (Brainard, 1997) running on Matlab (v2017a, MathWorks, MA, USA) was used to program all experiments. Some experimental tasks (Experiment 1 & 4-6) have previously been detailed (Bartoli et al., 2019) and are summarized below.

### Experiment 1: Perception task (visual categories)

In the perception task, all subjects were presented grayscale images from 10 visual categories (faces, houses, bodies, limbs, cars, words, numbers, instruments, corridors and phase-scrambled noise) in random order (see Figure 2a). Visual stimuli were from a publicly available corpus, previously used as a visual category localizer in human neuroimaging studies (Stigliani et al., 2015). On each trial, stimuli were shown for 1000 ms, with a random ISI between 1000 - 1500 ms. During the task subjects were required to press a button whenever they detected a specific stimulus being repeated back to back (1-back task). Performance was monitored by an experimenter present in the patient room. A total of 15 different stimuli were presented for each category, with 10 random images being repeated (serving as targets), leading to a total of 160 trials. On average the task was 7 minutes in duration.

### Experiment 2: Memory task (word-picture paired associates)

In the memory task, all subjects performed a paired-associates paradigm with an encoding and retrieval phase. During the encoding phase, subjects were presented with words (e.g., ‘coffee’, ‘nickel’) displayed above a box frame containing color photographs of well-known people (e.g., ‘Tom Cruise’ or ‘Julia Roberts’). Word stimuli are selected with limited letter range (4-8), number of syllables (1-3), concreteness rating (600-700) and imaginability rating (600-700) from the Medical Research Council Psycholinguistic Database (http://www.psy.uwa.edu.au/MRCDataBase/uwa_mrc.htm). Word-Picture pairs were presented for 5000 ms, with a self-paced ISI. Subjects pressed a button to advance to the next word-picture pair. A total of 15 word-picture pairs were presented for each task run. After a short delay, the retrieval phase would begin with subjects being presented with only cue words displayed above the box frame, but no picture. Cue words were from a list of 15 old (from encoding phase) and 15 new words. On each trial of the retrieval phase, subjects were required to retrieve the picture associated with the cue. Cue words were presented for 5000 ms and followed by a response period where subjects were asked to provide their memory strength of the cue and associate (see Figure 2). On average the task was 10 minutes in duration.

### Experiment 3: Resting task (eyes open / closed)

In the resting task, subjects (S 10,12,16) performed a simple task of alternating eyes open and closed periods, each lasting for 10 seconds. During the eyes open phase, subjects fixated on a central cross before being prompted to ‘close eyes’. During the eyes closed phase, subjects would keep their eyes shut until a brief auditory tone was played to signal the return to eyes open fixation. On average the task was 6 minutes in duration.

### Experiments 4-6: Supplemental perception tasks

To further examine our observations of comparable ripple attributes across states, particularly during perceptual tasks, in one subject (S12) we studied three additional perceptual tasks along with examining sleep (below). Details of these tasks have previously been reported (Bartoli et al., 2019) and are briefly summarized here. In the perception-grating task, full screen static grating stimuli are presented for 500 ms at 3 contrast levels (20, 50 & 100%). In the perception-color task, full screen color stimuli (9 colors) are presented for 500 ms. Finally, in the perception-color-object task, color images of different objects (Kiani et al., 2007) were presented for 500 ms.

### Sleep recording

Sleep recordings from subject S12 were also examined to provide an empirical within subject comparison of ripple properties across states. Sleep data was recorded overnight, during which polysomnography (PSG) recordings were performed to allow sleep staging. Following prior methods (Iber and American Academy of Sleep Medicine., 2007; Staresina et al., 2015; Ngo et al., 2020), PSG data was classified into awake, rapid-eye movement (REM) and non-REM (NREM) sleep states. We recorded 736 minutes (12.27 hours) of PSG data where 203 minutes (27.6%) were identified as awake, 78 minutes (10.6%) were identified as REM sleep, and 413 minutes (56.1%) were classified as NREM sleep. The remaining 42 minutes (5.7%) of recording were unable to be classified due to artifacts, and thus excluded from data analysis.

### Electrophysiological recording

Intracranial sEEG data was acquired at a sample rate of 2kHz and bandpass of 0.3-500Hz (4^th^ order Butterworth filter) using a BlackRock Cerebus system (BlackRock Microsystems, UT, USA). Initial recordings were referenced to a selected depth electrode contact within the white matter, distant from gray matter or pathological zones. During recordings, stimulus presentation was tracked using a photodiode sensor (attached to monitor) synchronously recorded at 30kHz. All additional data processing was performed offline.

### Data analysis: Preprocessing and spectral decomposition

All signal processing was performed using custom scripts in Matlab (v2018b, MathWorks, MA, USA). First, raw EEG signals were inspected for line noise, recording artifacts or interictal epileptic spikes. Electrodes with clear epileptic or artefactual activity were excluded from further analysis. Second, we identified sEEG probes targeting the medial temporal lobe, identifying which probes had electrode contacts within or at the boundary of the hippocampus. For each identified probe, hippocampal electrode contacts were notch filtered (60 Hz and harmonics) and rereferenced to a proximal electrode contact within the white matter on the same probe. Finally, Rereferenced signals from each hippocampal electrode were down sampled to 1kHz and spectrally decomposed using Morlet wavelets, with center frequencies spaced linearly from 2 to 200 Hz in 1 Hz steps (7 cycles).

### Data analysis: Ripple detection and rejection

After pre-processing steps, ripple events were identified in three general stages: i) time domain detection for identifying putative ripple events; ii) frequency domain assessment for accepting/rejecting ripple events; iii) electrode wise ripple count thresholding for inclusion/exclusion. Analytic criteria were based on prior human hippocampal ripples studies (Staresina et al., 2015; Helfrich et al., 2019; Jiang et al., 2019; Jiang et al., 2020; Ngo et al., 2020), as detailed below.

Signals from identified hippocampal electrodes (i.e., continuous voltage time-series filtered and re-referenced) were first band-pass filtered from 80 to 120 Hz (ripple band) using a 4^th^ order FIR filter. This ripple band range was selected based on prior studies in the human hippocampus (Bragin et al., 1999; Axmacher et al., 2008; Helfrich et al., 2019; Jiang et al., 2019; Norman et al., 2019; Vaz et al., 2019; Ngo et al., 2020), which report ripple range activity at lower frequencies than observed in the rodent (Buzsaki, 2015). Next, the root mean square (RMS) of the band-passed signal was calculated and smoothed using a 20-ms window. Ripples were detected based on amplitude and duration thresholds of this RMS time course. Whereby, ripple events were identified as having an RMS amplitude above 2.5, but no greater than 9, standard deviations from the mean. Ripple duration was defined as the supra-threshold time of the RMS-signal. Detected ripple events with a duration shorter than 38 ms (corresponding to 3 cycles at 80 Hz) or longer than 500 ms, were rejected. In addition to the amplitude and duration criteria the spectral features of each detected ripple event were examined. To do so, spectrally decomposed data (i.e. continuous time-series notch filtered, re-referenced and decomposed using Morlet wavelets), was used to calculated the frequency spectrum for each detected ripple event by averaging the normalized instantaneous amplitude between the onset and offset of the ripple event for the frequency range of 2-200 Hz. Spectral amplitude was normalized to a percent change signal by applying a baseline correction at each frequency based on the mean amplitude of the entire recording for a given electrode and frequency. For each detected ripple frequency spectrum, we examined the number and properties of spectral peaks. Spectral peaks were identified using the *findpeaks.m* MATLAB function. For every peak detected, the height, prominence, peak frequency and peak width were employed for our rejection evaluation. The impetus for these steps was to ensure that detected ripple events reflected high-frequency narrowband bursts limited to the ripple band range, rather than more broadband spectral changes increasing amplitude within and beyond the ripple band range (see Figure 1), driven by recording of ictal transient events in the time domain voltage. To do so we applied three criteria: First, ripple-events with a maximal spectral amplitude increase greater than 600% from the baseline were rejected. Second, genuine ripple events should display a unitary and predominant spectral peak with the ripple band range. Therefore, if no single prominent peak (ripple-peak) within the ripple band was identified, the event was rejected. Lastly, genuine ripple events should display a limited narrowband burst; therefore, if the ripple-peak had a wide peak width (more broadband spectral change) or a prominent high frequency activity (large peaks in 120 – 200 Hz), the event was rejected.

**Figure 1.**
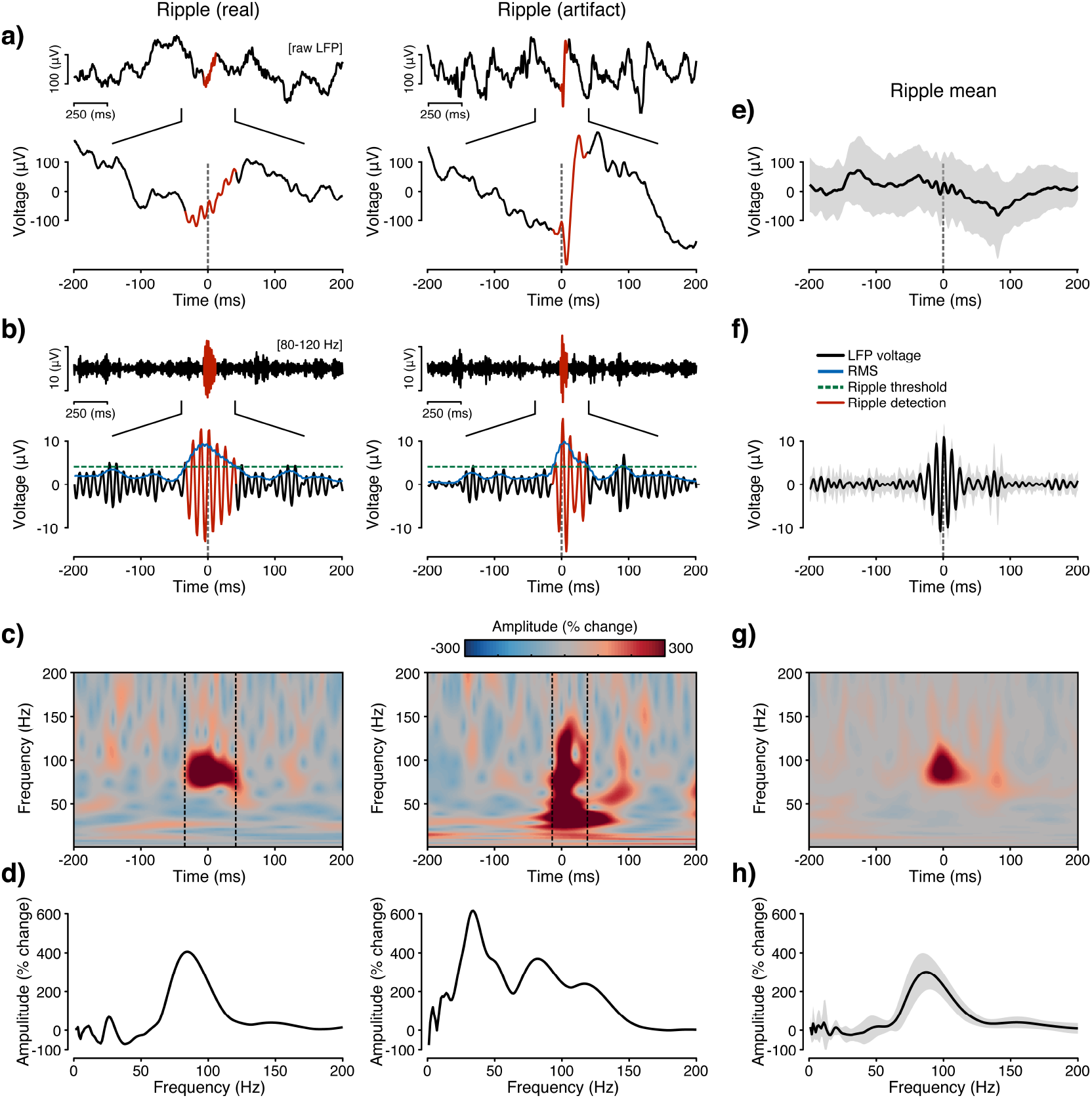
Ripple detection and artifact rejection. **a)** Example raw voltage traces showing detected ripples that are subsequently identified as real (left) or artifactual (right; data from S10, electrode 4). Here and below, time zero aligns to the maximal ripple amplitude, with detected ripple events shown in red. **b)** Ripple-band (80–120 Hz) voltage traces for example real and artifactual ripples (same data as (a)). Green dashed line reflects the ripple detection threshold for this electrode. Blue line reflects the rippleband RMS envelope. **c)** Spectrograms are shown for the example real and artifactual ripples, color maps reflect percentage change in amplitude relative to the total signal mean. Black dashed lines denote the onset and offset of detected ripples. Real ripple events display a high-frequency narrow band time-frequency representation, while the artifactual ripple displays a much more broadband frequency representation due to the sharp voltage transient shown in (a). **d)** Normalized amplitude spectra (percentage change) for the example real and artifactual ripples averaged over the detected ripple onset/offset window. While the real ripple displays a predominant spectral peak in the ripple-band range, the artifactual ripple shows multiple spectral peaks outside of the ripple-band. In order to isolate genuine ripple events, these spectral features where quantified and incorporated into our ripple detection algorithm (see methods). **e**) Mean ripple triggered raw voltage trace after artifact rejection (ripple real/artifact = 10/17; here and below error shading reflects s.e.m). **f–h)** Same plots as (b-d) but for the mean ripple triggered data: ripple-band voltage (f), spectrogram (g) and normalized amplitude spectra (h), respectively. By identifying and removing artifactual ripples, mean ripple data displays a clear ripple signature that is spectrally peaked and isolated in the ripple-band.

To quantify these criteria, we implemented the following steps for rejection 1) events with more than one peak in the ripple band; 2) events where the most prominent and highest peak was outside the ripple band and sharp wave band for a frequencies >30 Hz (as peaks found below 30 Hz may reflect the sharp wave that usually accompanies ripple events (Jiang et al., 2020)); 3) events where ripple-peak width was greater than 3 standard deviations from the mean ripple-peak width calculated for a given electrode and recording session; and 4) events where high frequency activity peaks exceed 80% of the ripple peak height. After applying the trial-wise spectral rejection, we calculated the number of ripples detected and ripple rejection rate for each electrode. We then rejected any electrode with a low ripple count (<20 ripples detected per electrode per task) or high rejection rate (greater than 30 % rejection rate), as these likely reflect weak or noisy recordings.

For identified ripples, there were four main attributes we examined: 1) ripple rate (Hz) calculated by using the number of detected ripple events divided by recording time in seconds, 2) ripple duration (ms), 3) ripple max amplitude (uV) calculated using ripple band-passed signal, and 4) ripple peak frequency (Hz).

### Statistical Analysis

Ripple event data was subject to quantification using parametric and non-parametric statistical methods as appropriate for underlying data distributions. Mixed-effects models, using subject and electrode as random effects, were employed to account for the nesting of multiple electrodes within each subject. Statistical analyses were carried out using R statistical software (R Development Core Team, 2010).

## Results

### Detected ripple events

Direct hippocampal recordings were performed in 17 subjects undergoing invasive monitoring for the surgical treatment of refractory epilepsy (see methods). Across subjects, a total of 127 electrodes were anatomically localized to the hippocampi. During recordings, subjects performed three tasks focused on visual object perception (Perception), episodic memory encoding and retrieval (Memory), and eyes open/closed rest (Resting; additional supporting tasks described below). Using threshold-based ripple detection across these tasks, we identified a total of 13,520 ripple events, with 77.4% of these detected ripples subsequently being classified as genuine (10,465/13,520; see Figure 1 for examples of real and artifactual ripple events). In addition, after applying an electrode-rejection criterion, 82 electrodes were kept for our main analysis, resulting in a total of 7,831 ripples (mean ripple keep rate per electrode = 86.27%). See methods for ripple detection and rejection details. Next, we sought to quantify the attributes of hippocampal ripple activity across cognitive tasks.

### Ripple attributes across different cognitive tasks

After applying time and frequency domain metrics for hippocampal ripple identification, we observed clear ripple events in all subjects and across all tasks. First, we sought to quantify ripple attributes across our three main tasks (Perception, Memory and Resting, Figure 2). As shown in Figure 2, hippocampal ripples were readily detected across all three tasks, displaying highly similar electrographic signatures (Figure 2e-g). Given these similarities, we also quantified several other ripple attributes for comparison across tasks, which included: i) rate (Hz, number of ripples per second); ii) duration (ms, time period ripple is above amplitude threshold); iii) amplitude (μV, max ripple amplitude) and iv) peak frequency (Hz, frequency with max amplitude within ripple band). Qualitatively, ripple attributes did not display any large differences across tasks. However, group linear mixed-effects analysis modeling the ripple attributes with tasks as a fixed effect, and treating subject and electrode as random effects, and Satterthewaite approximations to test the significance of the model coefficients, revealed a significant main effect of task for rate, duration, and amplitude but not for peak frequency. Specifically, using pairwise Tukey’s range test, with p-values adjusted for comparing a family of three estimates, ripple rates were higher for Resting than the other two tasks [Resting - Memory: *t*(112) = 3.354, p = 0.0031 and Resting - Perception: *t*(112) = 2.522, p = 0.0348]; ripple duration was longer in Resting than other two tasks [R-M: *t*(119) = 4.942, p <0.0001 and R-P: *t*(119) = 4.659, p <0.001]; and ripple amplitude was greater in Resting than other two tasks [R-M: *t*(109) = 3.406, p =0.0026 and R-P: *t*(109) = 2.192, p = 0.001]. Moreover, there was no significant difference between Memory and Perception tasks for any of the ripple attributes. These results suggest that ripple attributes are quite comparable between the two active cognitive tasks (Perception and Memory) and modestly different from the offline resting state, which shows slightly more pronounced ripple activity. Given the general stability of ripples across tasks, we next examined event-related changes in ripple rates, given prior evidence of task modulation of ripple activity (Axmacher et al., 2008; Norman et al., 2019; Vaz et al., 2019).

**Figure 2.**
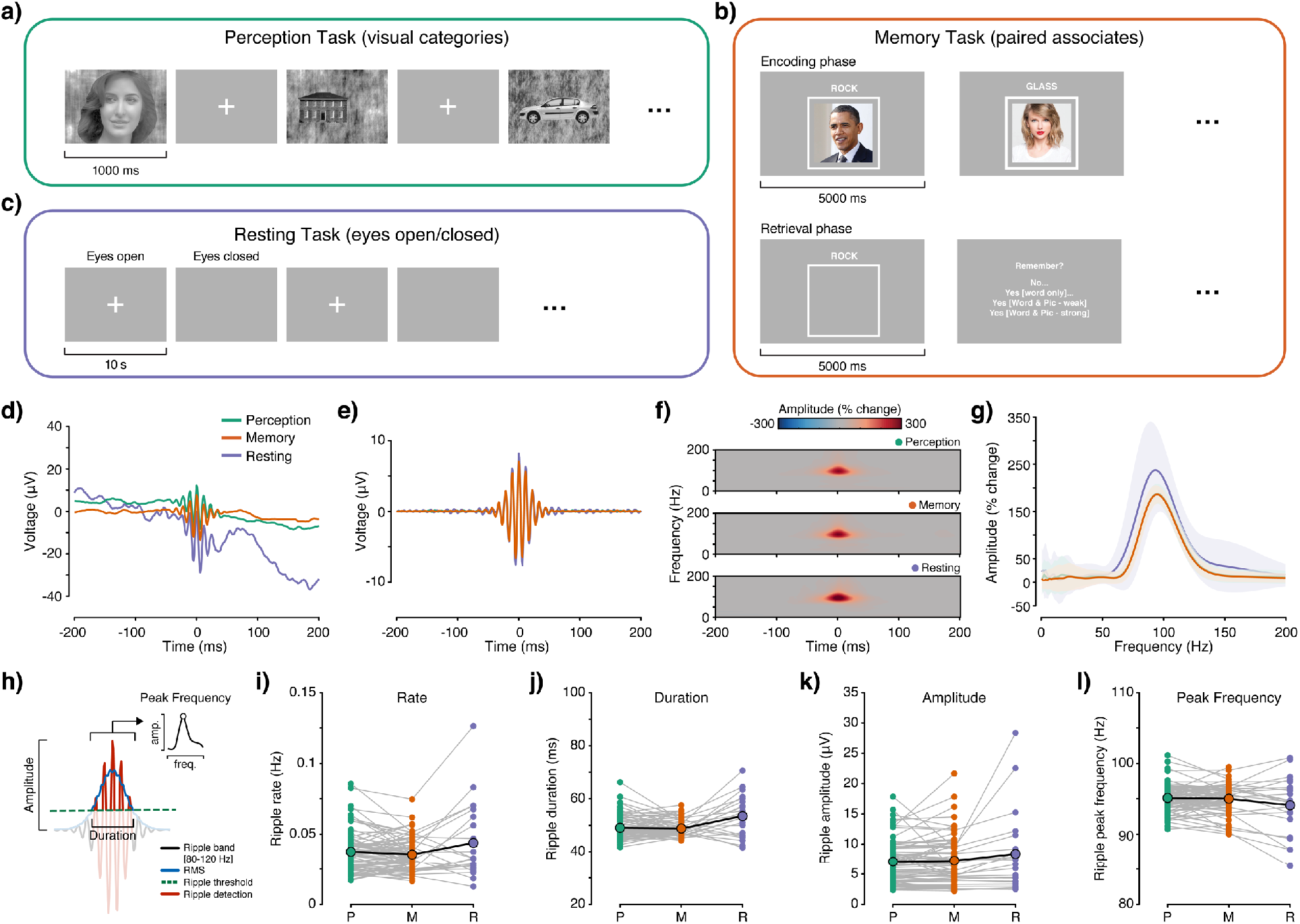
Ripple attributes across different cognitive tasks. **a)** Perception task stimuli and experimental procedure. Grayscale images from ten visual categories were presented for 1000 ms with a random ISI of 1-1.5s. Subjects had to provide a button press to indicate any 1-back stimuli repetitions. **b)** Memory stimuli and experimental procedure. During the encoding phase, word-image pairs were presented for 5000 ms with a self-pace ISI. During the retrieval phase, cue words were presented for 5000 ms followed by a memory strength judgement. Subjects were required to encode word-face associations, and to retrieve the associated face image from presented word cues (which included new cue words). **c)** Resting task experimental procedure. During the resting task, subjects alternated between 10 second periods of eyes-open and -closed based on visual or auditory cues. **d)** Group averaged ripple-triggered voltage trace for each task. **e)** Group averaged ripple-band triggered voltage trace for each task. **f)** Group averaged ripple-triggered spectrograms for each task, color maps reflect percentage change in amplitude relative to the total signal mean. **g)** Group averaged normalized amplitude spectra (percent change) for each task averaged over the detected ripple onset/offset window. Overall, hippocampal ripples were reliably detected across all three tasks, with highly comparable spectro-temporal properties. **h)** Schematic of ripple attribute quantification. **i-l)** Ripple attributes are shown for all subjects and electrodes across tasks: rate (i); duration (j), amplitude (k) and peak frequency (l). Statistical analysis revealed a significant main effect of task for rate, duration and amplitude, where these ripple attributes were significantly greater for the Resting compared with the Memory and Perception tasks.

### Event-related ripples across tasks

To quantify even-related changes in ripple rates across tasks, detected ripple events were aligned to stimulus presentation conditions across all tasks (Perception, Memory & Rest). In addition, data from the Memory task was separated into encoding/retrieval phases, while the Rest task was separated into eyes open/closed phases. Event-related ripple events are shown for all tasks, subjects, electrodes and trials in Figure 3. As noted above, and consistent with prior work, all tasks show clear evidence for ongoing ripple events. However, when considering event-related changes in ripple rates, no overt modulation was observed across tasks. This was true also when considering relative changes in ripple rate, where the ripple rate time course was normalized by subtracting the mean ripple rate during the pre-stimulus baseline (500 ms) (Figure 3c,g,k). While more modest in sample size, ripple rates were significantly higher for eye closed, compared with eyes open (Wilcoxon rank sum test, W = 52513, p <0.0001), in the Resting task. Behaviorally, we observed high accuracy for the 1-back Perception task (mean hit rate 75.85 %) and Memory retrieval task (mean hit rate 81.57% and mean correct rejection rate 84.25%). Non-parametric spearman correlation revealed no significant relationship between mean ripple rate and task accuracy (Perception task: ρ = 0.25, p = 0.42; Memory task: hit rate ρ = 0.32, p = 0.21 & correct rejection rate ρ = 0.12, p = 0.64). Overall, these data are consistent with the observations noted above, that hippocampal ripples occur at a stable rate during cognitive tasks, showing minimal task modulation and more sensitivity to general state changes (e.g. online vs. offline states). Therefore, we next examine addition factors which may influence ripple events.

**Figure 3.**
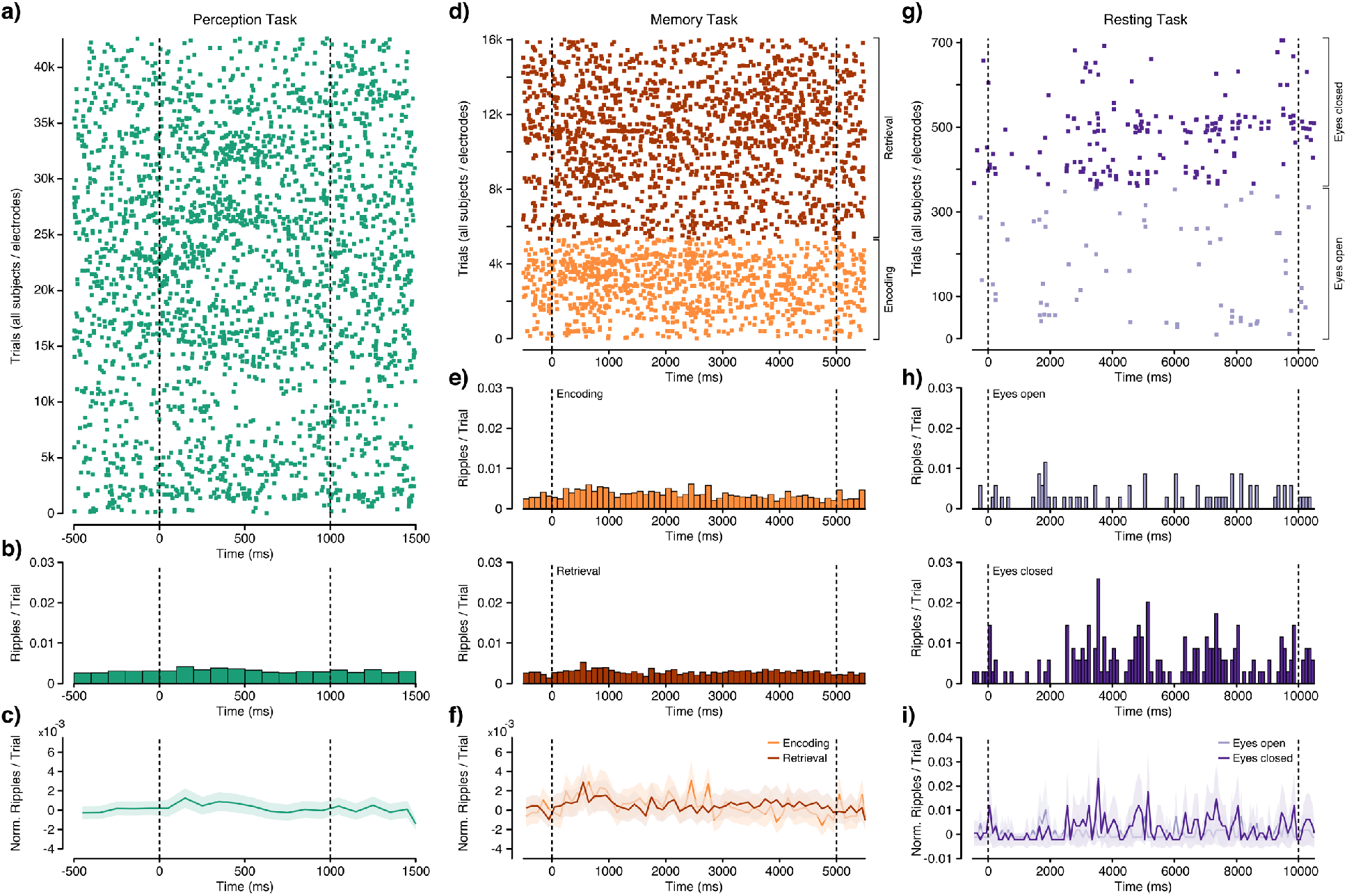
Event-related ripples across tasks. Event-related ripple events across tasks for all subjects, electrodes and trials. **a)** Ripple events for the Perception task (all electrodes x trials, n = 42,582), green markers indicate a single identified ripple event. Dashed lines indicate stimulus presentation period. **b)** Mean ripple rate across trials. **c)** Pre-stimulus baseline corrected ripple rate across trials, error shading reflects 3 s.e.m. **d)** Ripple events for the Memory task (all electrodes x trials, n = 16,065), markers indicate a single ripple event for encoding (orange) and retrieval (brown) trials. Dashed lines indicate stimulus presentation period. **e)** Mean ripple rates for encoding and retrieval trials. **f)** Pre-stimulus baseline corrected ripple rates for encoding and retrieval, error shading reflects 3 s.e.m. **g)** Ripple events for the Rest task (all electrodes x trials, n = 708), markers indicate a single ripple event for eyes open (light purple) and eyes closed (purple) trials. Dashed lines indicate start and end of the eyes open/closed period. **h)** Mean ripple rates for eyes open and eye closed trials. **i)** Pre-stimulus baseline corrected ripple rates for eyes open and eyes closed, error shading reflects 3 s.e.m.

### Ripple attributes across hemisphere and time

To assess whether ripple attributes (rate, duration, amplitude and peak frequency) were affected by other recording variables, we examined these attributes across recording sites, number of days post implant surgery, and the time of day tasks were conducted (Figure 4). Consideration of these factors are important given the potential confound of pathophysiological activities, and for exploring any putative functional impacts of behavior. First, we compared ripple attributes between left and right hemispheres. Of the 82 included electrodes (Figure 4a for the anatomical location), 49 electrodes were identified in left hippocampus. Similar to the group results, ripple attributes did not show any large difference between left and right hemisphere (Figure 4b-e). Nonparametric Wilcoxon rank sum test comparing ripple attributes between left and right hemisphere revealed no significant difference in rates (W = 629, p = 0.08), duration (W = 722, p = 0.38) and peak frequency (W = 676, p = 0.19); and only marginally significant effects in amplitude (W = 1030, p = 0.04). Next, we examined the relationships between ripple attributes and days post implant surgery. Most task recordings were performed between 1-5 days post implantation, with ripple attributes being stable across days (Figure 4f-i). Non-parametric spearman correlation revealed no significant correlation between days post implant and any of the ripple attributes (rate: ρ = −0.05, p = 0.27; duration: ρ = − 0.09, p = 0.06; amplitude: ρ = 0.01, p = 0.80; frequency: ρ = 0.07, p = 0.16). Lastly, we examined the relationship between ripple attributes and the time of day recordings took place. Recordings were performed between 10 am to 7 pm, with the ripple attributes being stable throughout the day (Figure 4j-m). Non-parametric spearman correlation revealed no significant correlation between time of the day with any of the ripple attributes (rate: ρ = 0.074, p = 0.11; duration: ρ =0.07, p =0.14; amplitude: ρ =0.02, p = 0.59; frequency: ρ = −0.09, p = 0.05). These results suggest that hippocampal ripples, and their attributes, do not differ between hemisphere and are stable across and within days. Such findings suggest a limited impact of pathophysiological factors impacting overall ripple statistics. In turn, they also suggest ripples occur at a low and regular rate throughout the waking day and do not appear to show any pronounced modulation specific to memory behavior. Consistent with a large literature, ripple rates are expected to be more pronounced during offline states. While we observed modest evidence for this during the Resting task, offline sleep states have shown the most pronounced effects. In addition, we only examined one perceptual task which included face, scenes and objects not unlike our memory paradigm. Therefore, in an attempt to empirically consider these factors, we examined ripple attributes across additional perception tasks as well as during sleep.

**Figure 4.**
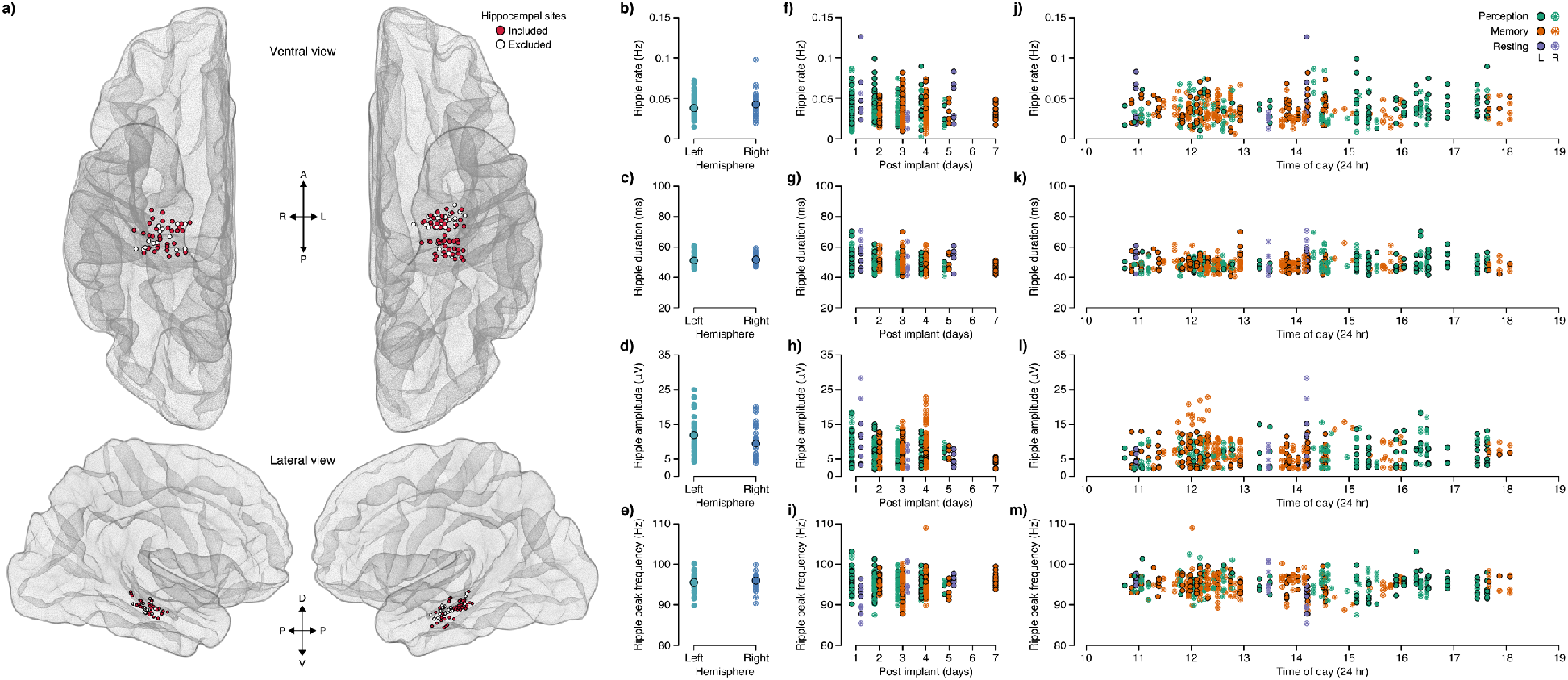
Ripple attributes across hemisphere and time. **a)** Anatomical location of identified hippocampal electrodes from all subjects, normalized to MNI space. Included electrodes, which passed a selection criterion, are shown in red with excluded electrodes shown in white (see methods). Ripple attributes of rate, duration, amplitude and peak frequency are shown as a function of anatomy and time. **b-e)** Ripple attributes (rate, duration, amplitude, peak frequency) are shown for all electrodes, averaged across tasks, from the left and right hemispheres. **f-i)** Ripple attributes (rate, duration, amplitude, peak frequency) are shown for each electrode, task (perception, memory, resting) and hemisphere (left/right) as a function of the days post electrode implantation. **j-m)** Ripple attributes (rate, duration, amplitude, peak frequency) are shown for each electrode, task (Perception, Memory, Resting) and hemisphere (left/right) as a function of the time of day recordings were performed. Overall, hippocampal ripple attributes were similar between hemispheres and stable across and within days.

### Ripple attributes across multiple task and state conditions

In one subject (S12), we carried out three additional perception tasks (Perception – grating, Perception – color and Perception – color obj.), as well as recording one night of sleep to assess how task and state conditions influenced ripple attributes. We sought to examine 1) whether ripple attributes were comparable amongst cognitive tasks and 2) whether ripple attributes during sleep, a pronounced offline state, differed from other cognitive states. As shown in Figure 5, hippocampal ripples were readily detected across all six tasks and three sleep stages, displaying highly similar electrographic signatures with the only exception for NREM sleep (Figure 5b-e). Notably, in the subject averaged ripple-triggered raw voltage trace plot (Figure 5b), ripples recorded during NREM sleep were nested in a sharp wave that displayed large voltage deviation compared to other cognitive tasks and sleep stages. In addition, the normalized amplitude spectra for NREM sleep ripples (Figure 5e) showed a higher amplitude percent change, but lower frequency, ripple band peak compared to other cognitive tasks and sleep stages. These observations illustrate a clear difference between NREM sleep ripples and other tasks and sleep stage ripples. Therefore, to compare across tasks and sleep stages, we carried out linear mixed-effects analysis modeling ripple attributes with task/sleep stage as a fixed effect and treating electrode as random effect. Satterthewaite approximations to test the significance of the model coefficients revealed a significant main of task/sleep stage for rate, duration, amplitude and peak frequency. Using pairwise Tukey’s range test, p-value adjusted for comparing a family of 9 estimates (Perception, Memory, Resting, Perception – grating, Perception – color, Perception – color obj., Awake, REM sleep and NREM sleep), we compared the ripple attributes amongst tasks and sleep stages and summarized the results in Table 2. Specifically, ripple rate was significantly higher in NREM sleep than other tasks and sleep stages; ripple duration was significantly longer in NREM sleep than Memory, Awake, and REM sleep; ripple amplitude was significantly greater in NREM sleep than other tasks and REM sleep, but not in Awake; and lastly, ripple peak frequency in NREM sleep was significantly lower than other tasks and sleep stages, except in Perception-grating. Overall, ripple attributes during NREM sleep were distinctly different from other cognitive states and sleep stages.

**Figure 5.**
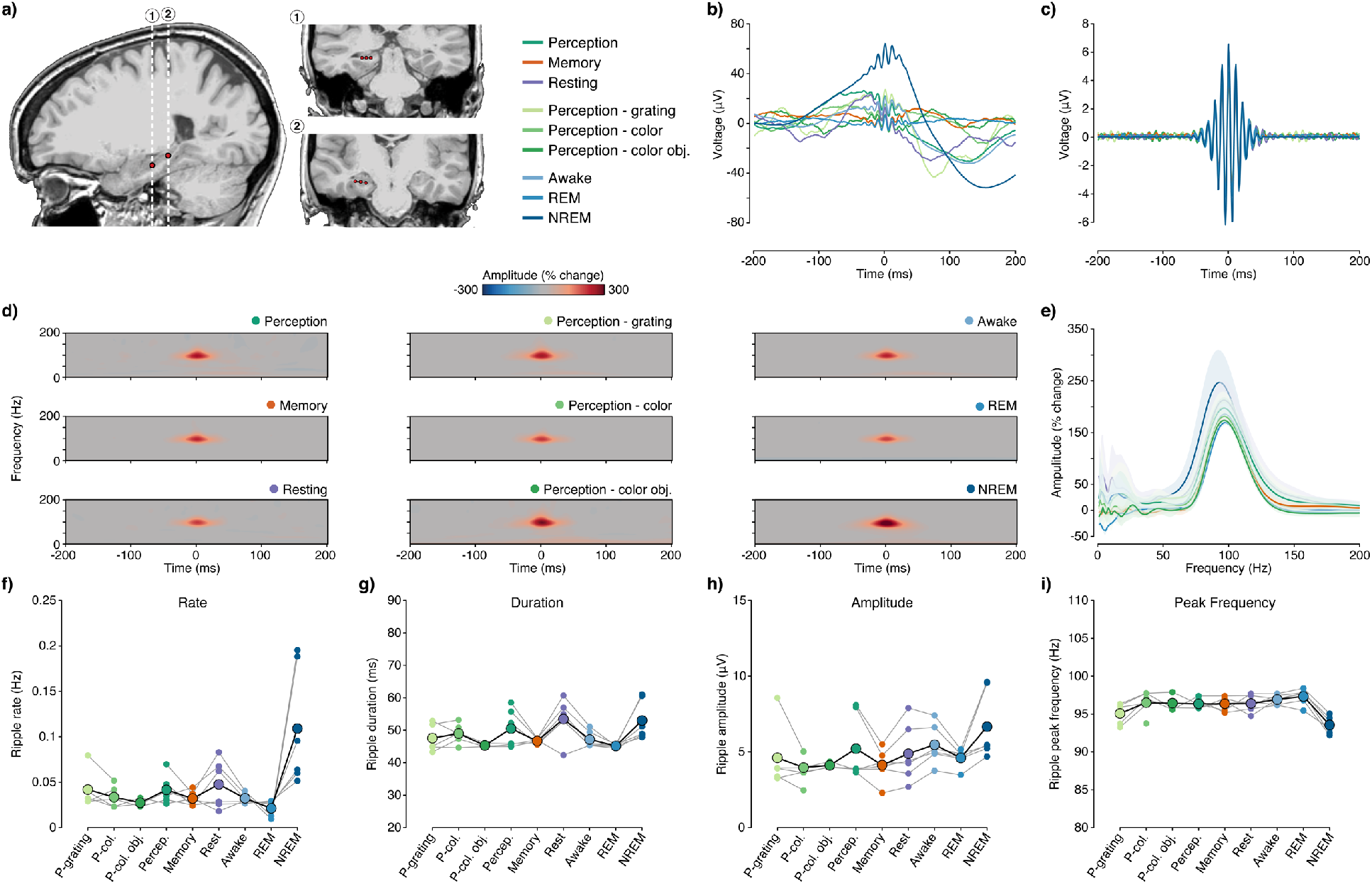
Ripple attributes across task and sleep states. **a)** Anatomical Location of hippocampal electrodes (n = 6) in subject S12 for two sEEG depth probes. Sagittal view (left) shows longitudinal position of each probe, with the white dashed lines indicated the respective coronal slices (right; *1* and *2*). **b)** Subject averaged ripple-triggered raw voltage trace for each task and sleep stage. **c)** Subject averaged ripple-triggered voltage trace within the ripple band for each task and sleep stage. **d)** Subject averaged ripple-triggered spectrograms for each task and sleep stage, color maps reflect percentage change in amplitude relative to the total signal mean. **e)** Subject averaged normalized amplitude spectra (percent change) for each task and sleep stage, estimated over the detected ripple onset/offset window. Overall, hippocampal ripples were reliably detected across all six tasks and three sleep stages, with highly comparable spectro-temporal properties. However, it is noticeable, that ripple-trigger activity during NREM is uniquely nested within a slower frequency oscillation, as previously observed [refs]. **f-i)** Ripple attributes are shown for all electrodes across tasks and sleep stages: rate (f), duration (g), amplitude (h) and peak frequency (i) (small circles reflect electrodes, large circles reflect mean). For some tasks and electrodes no ripples were detected. Statistical analysis revealed a significant main effect of task and sleep stage, whereby ripple attributes recorded during NREM were significantly different from the other tasks and sleep stages (see results).

**Table 2.** Pairwise Tukey’s range test comparing ripple attributes: rate, duration, amplitude and peak frequency between NREM sleep and cognitive tasks and other sleep stages, p-value is adjusted for comparing a family of 9 estimates.

| **Tasks** | **Ripple attributes** |  |  |  |
| --- | --- | --- | --- | --- |
|  | *Rate* | *Duration* | *Amplitude* | *Frequency* |
| *NREM –* |  |  |  |  |
| Perception | $t(43.8) = 5.592$,<br>$p < 0.0001$ | $t(43.9) = 1.578$,<br>$p = 0.8110$ | $t(44) = 3.472$,<br>$p = 0.0291$ | $t(43.8) = -5.817$,<br>$p < 0.0001$ |
| Memory | $t(43.8) = 6.380$,<br>$p < 0.0001$ | $t(43.8) = 3.956$,<br>$p = 0.0077$ | $t(44) = 6.044$,<br>$p < 0.0001$ | $t(43.8) = -5.832$,<br>$p < 0.0001$ |
| Resting | $t(43.8) = 5.098$,<br>$p = 0.0002$ | $t(43.9) = -0.272$,<br>$p = 1.000$ | $t(44) = 4.248$,<br>$p = 0.0032$ | $t(43.8) = -5.908$,<br>$p < 0.0001$ |
| P-grating | $t(44.1) = 5.129$,<br>$p = 0.0002$ | $t(44) = 2.878$,<br>$p = 0.1224$ | $t(44) = 3.996$,<br>$p = 0.0068$ | $t(44.1) = -2.775$,<br>$p = 0.1588$ |
| P-color | $t(43.8) = 6.260$,<br>$p < 0.0001$ | $t(43.9) = 2.514$,<br>$p = 0.2533$ | $t(44) = 6.442$,<br>$p < 0.0001$ | $t(43.8) = -6.105$,<br>$p < 0.0001$ |
| P-color obj | $t(44.7) = 4.802$,<br>$p = 0.0006$ | $t(44.4) = 2.735$,<br>$p = 0.1650$ | $t(44.1) = 3.654$,<br>$p = 0.0179$ | $t(44.6) = -4.802$,<br>$p = 0.0057$ |
| Awake | $t(43.8) = 6.366$,<br>$p < 0.0001$ | $t(43.9) = 3.672$,<br>$p = 0.0170$ | $t(44) = 2.835$,<br>$p = 0.1341$ | $t(43.8) = -7.010$,<br>$p < 0.0001$ |
| REM | $t(43.8) = 7.283$,<br>$p < 0.0001$ | $t(43.9) = 4.917$,<br>$p = 0.0004$ | $t(44) = 4.921$,<br>$p = 0.0004$ | $t(43.8) = -7.854$,<br>$p < 0.0001$ |

## Discussion

Using direct intracranial recordings from the human hippocampus, we quantified hippocampal ripple attributes across different cognitive tasks, sleep stages, recording hemispheres, and potential pathophysiological factors. Our results highlighted the stability of ripples attributes across cognitive tasks. Ripple rate, duration, amplitude, and frequency were comparable between Perception and Memory tasks; however, they differed from offline resting conditions, most notably NREM sleep. Moreover, ripples attributes were stable between recording hemispheres, throughout the time of day and proximity to electrode implantation date.

Hippocampal ripples are short bursts of high-frequency activity (80 – 120 Hz in humans) that have been linked to memory consolidation, specifically the reactivation or replay of memory content during offline states such as NREM sleep (Diekelmann and Born, 2010; Joo and Frank, 2018; Klinzing et al., 2019). More recently, hippocampal ripples have been recorded during awake online states, specifically during episodic memory tasks (Norman et al., 2019; Vaz et al., 2019). While it is important to examine the functional significance of ripples in facilitating memory encoding and retrieval, it is unclear if ripples occur in other cognitive tasks or how the properties of ripple differ between cognitive states or task demands. Work in rodents exploring the differences across brain states has demonstrated that the occurrence of ripples is much lower during awake compared to prolonged immobility or sleep (Buzsaki, 2015; Joo and Frank, 2018). Furthermore, ripple attributes such as amplitude, duration, and peak frequency have been reported to differ between awake exploration and sleep and rest (Buzsaki, 2015), which could be important neural markers for cognitive processes. Here, we examined ripples attributes across our three main tasks, two cognitive tasks, and offline resting (Figure 2). These tasks were chosen so as to directly compare ripples found during episodic-memory (Memory) vs non-episodic-memory tasks (Perception). We also included an offline awake resting condition for comparing online and offline brain states. In general, we found that ripple attributes were stable between awake cognitive tasks regardless of the engagement in episodic memory behavior. However, during resting states, ripples showed a higher occurrence rate, were longer in duration and larger in amplitude. These data suggest that hippocampal ripples are not uniquely related to episodic-memory behavior, given that ripple attributes were comparable between episodic and non-episodic tasks. As noted above, the finding of enhanced ripple activity during offline states is consistent with observations in the rodent and human literature (Brazdil et al., 2015; Joo and Frank, 2018). The largest difference in ripple attributes were observed during NREM sleep where ripple rate was higher, duration longer, amplitude greater but peak frequency slower in contrast to any other cognitive task or sleep stages. This result is in line with the idea that hippocampal ripples play a pivotal role in NREM sleep mediated memory consolidation (Klinzing et al., 2019).

While we reliably detected hippocampal ripples across all tasks, some aspects of ripple attributes differ from prior work. Firstly, ripple rates detected during cognitive tasks were much lower compared to prior work. For example, recent work by Vaz et al., 2019 reported a mean ripple rate of 0.21 ± 0.02 Hz across participants during an episodic task, and soon after Norman et al. 2019 reported ripple rates of 0.41 Hz across all experimental conditions; in contrast our averaged ripple rate across the three main tasks was an order of magnitude smaller at 0.037 ± 0.013 Hz. Secondly, prior studies have highlighted increased ripple rates during encoding and retrieval were indicative of successful memory (Norman et al., 2019; Vaz et al., 2019). We did not observe any clear event-related change in ripple rate in the Memory or Perception task, although difference between eyes-open and closed were observed for the Resting task.

Several factors may account for these differences. One important consideration is the anatomical locations of ripple recording sites. For example, Vaz et al. (2019) examined ripple events from the medial temporal lobe (MTL) cortical structures including entorhinal cortex and parahippocampal gyrus, but not the hippocampus. Consequently, higher ripple rates may be present in neocortical structures, although it is typically viewed that the hippocampus is the key generator of ripple activity (Axmacher et al., 2008; Buzsaki, 2015). However, Norman et al. (2019) recently reported similar findings which included direct hippocampal recordings, observing higher rates as noted above. Therefore, while ripple events have been historically viewed as a hippocampal phenomenon, they may be observed in the neocortex, and with higher rates or occurrence, but such factors do not fully account for differences with the present study.

Another critical factor is the ripple detection method employed. Given that direct hippocampal recordings in human subjects are collected from individuals undergoing invasive monitor as part of their treatment for refractory epilepsy, many studies have expressed concerns for reliable and genuine ripple detection (Jiang et al., 2020). Particularly, many electrodes localized in hippocampus are proximal to or within seizure onset zones, which generate high frequency oscillations (HFOs; 80 - 500 Hz) and inter-ictal epileptic spikes (Jacobs et al., 2012). Therefore, simple ripple detection methods may be prone to false positives (see Figure 1 for examples). In order to remedy this possible false detection, prior studies have employed different validation methods including visual inspection of randomly sampled data (Vaz et al., 2019), excluding simultaneously detected events on multiple channels, or proximity of ripple events occurrence to pathological events (Norman et al., 2019), as well as other steps for artefact rejection.

To reliably detect genuine hippocampal ripples, we applied additional time-frequency criteria to reject potential false positives after initial identification via ripple band threshold-based detection. First, we included a conservative duration rejection threshold of 38 ms which corresponds to three cycles at 80 Hz, the lower bound of the ripple band frequency. This duration threshold ensured that the detected signal is a sustained oscillatory event rather than a single isolated transient signal change such as an inter-ictal spike or recording noise. It is worth noting that Vaz et al. 2019 reported 34 ms as their average duration recorded from MTL electrodes with an estimated range of 25 to 60 ms. Therefore, our approach would have excluded some of the short duration ripples reported. However, it is important to point out that in a follow up analysis, Vaz et al. excluded short duration ripples (25 - 30 ms) and still found significant memory effects. Secondly, we included a ripple spectra rejection method where events that did not exhibit a stereotypical ripple spectra profile were excluded, consistent with some prior sleep studies (Jiang et al., 2019; Jiang et al., 2020; Ngo et al., 2020). This approach allowed us to look beyond the typical bandpass filtered ripple band data. Examining the wide band spectral change enabled us to identify low frequency noise, broadband transients or HFOs that could be misidentified as a ripple event. Together, these criteria likely account for the lower ripple event rates observed in the present study. Importantly, we note that this approach may be too conservative, such that our false negative rate maybe inflated. It is important to note that we also excluded electrodes if the spectral based rejection rate was >30% of all identified ripple events. This step aimed to both exclude noisy electrodes, not identified in early stages of processing, and the erroneous creation of low rate recordings. While establishing ground truth for ripples rates within the human hippocampus can be difficult, we believe our reported rates fall within an appropriate range. Central to this conclusion is that prior work in human hippocampus focused on NREM sleep, where ripples rates are thought to be maximal, typically report rates within the range of 0.1 – 0.5 Hz (Jiang et al., 2020). Additionally, a subset of these sleep focused studies also reported ripple rates during awake states with a range of 0.01 – 0.1 Hz (Jiang et al., 2020). These ripple rate ranges are more comparable to our observations. Indeed, as shown in Figure 5, when applying our methods to NREM sleep data we observer similar ripple rates, which importantly are on average larger than for task conditions. Moreover, our relatively low ripple occurrence was also reported by prior work in non-human primates. For example, Logothetis et al. (2012) examined hemodynamic neuroimaging responses time locked to electrophysiological hippocampal ripples events in the macaque. Using ripple band threshold detection as well as spectral selection, similar to the current study, these authors reported ripple rates of 5.6 per minute (0.09 Hz) during unanesthetized spontaneous recordings. In addition to similar rates of ripple events, ripple duration and peak frequency in the macaque were highly similar to our observations in the human (Logothetis et al., 2012). These low rates in the macaque may be related to a lack of task engagement, however, similarly low rates have been observed in the macaque hippocampus during perceptual memory tasks (Leonard et al., 2015; Leonard and Hoffman, 2017). For example, Leonard et al. observed low rate hippocampal ripple events during complex visual scene viewing (max rate ~1.12/minute, 0.019 Hz), which showed a slow drifting increase during visual search (Leonard et al., 2015) that was subsequently shown to be enhanced for repeated vs. novel scenes (Leonard and Hoffman, 2017). Such slow rates are similar to our lack of rapid event-related changes in ripple rates and suggest modulations in rate occur over slower timescales. This view is consistent with prior observations in the rodent showing a tight anti-correlated coupling between slow changes in arousal states (i.e. pupil dilation) and hippocampal ripple rate (McGinley et al., 2015a; McGinley et al., 2015b). In addition, it is consistent with ripple events occurring via intrinsic mechanisms, rather than externally driven stimulus events (Kay and Frank, 2019). Together, our findings replicate prior observations of online hippocampal ripple events, but differ in the mean rate of ripples and a lack of clear event related ripple modulation. While these differences may be related to the detection methods employed, our observations are consistent with prior sleep studies in humans, and other non-human primate studies examining hippocampal ripple activity (Logothetis et al., 2012; Leonard et al., 2015; Helfrich et al., 2019; Jiang et al., 2020; Ngo et al., 2020).

In addition to examining task demands and brain state modulation of ripple attributes, we also considered other factors that may influence ripple rate. We observed no statistically significant difference in ripple attributes between hemispheres, both in aggregate and when considering specific tasks. Also, we examined whether our observations were potentially biased by patient state factors. To do so we quantified ripple attributes as a function of the days post electrode implantation, and as a function of the time of day recordings were performed. However, we observed no correlation of ripple attributes with either chronological variable, for a reasonable spread of days post implantation (1-7 days) and time of day (~11am-6pm). These findings suggest hippocampal ripple attributes are stable throughout the day, making our comparisons across tasks unlikely to be confounded by these factors. More generally, these observations further support the notion of ripple events being stable across a host of conditions, and being most sensitive to overt behavioral state changes (e.g. online vs. offline).

Why might hippocampal ripples occur with such stability throughout waking states? One natural hypothesis would be that ongoing and sparse hippocampal ripple events aid in establishing latent memory traces, consistent with two-stage models of systems consolidation (Dudai et al., 2015; Kumaran et al., 2016; Klinzing et al., 2019). This view might lead to evidence that ripple events are modulated by factors associated with subsequent memory behavior, as previously observed, however was not strongly supported by our findings. It is important to note, that while subjects did perform an explicit episodic memory task, this is not a unique period of putative memory encoding, as we expect subjects to later recall performing other tasks as a simple consequence of the own autobiographical memory. It will therefore be important for future studies to carefully quantify key behavioral factors which influence hippocampal rates and the veracity of their neocortical counterparts. In doing so, continued convergence of evidence across species can be obtained to further elucidate the mechanisms of memory consolidation. Additionally, ripples have been identified as the most prominent self-organized event in hippocampus and may reflect the default pattern of hippocampal circuits. Therefore, changes in ripple attributes could be indicative of brain states (Buzsaki, 2015; Kay and Frank, 2019). This view might lead to evidence that ripple events are modulated by factors associated with states of the brain, such as arousal as noted above. The stable ripple rates we observed could be a result of overall similar arousal during these tasks. While we did not directly assess the arousal levels of our participants, the crude separation of ‘online’ and ‘offline’ tasks did reveal changes in ripple attributes associated with brain state.

In conclusion, our findings integrate and extend prior work in the human, non-human primate and rodent hippocampus to suggest that ripple events are relatively stable across cognitive tasks. Ripple events were not unique to episodic-memory behavior, occurring with similar attributes during other perceptual tasks, but less frequently than during ‘offline’ tasks and brain states. Such findings add growing support to the view that hippocampal ripples serve as an internally generated and state dependent mechanism for establishing, and subsequently strengthen, memory traces in the mammalian brain.

## Acknowledgments

The authors thank Hong-Viet Ngo and Bernhard Staresina for assisting with sleep staging neural data. This work was supported by NIH grants R01EY023336 to D.Y., R01MH106700 to S.A.S., and R01MH116914 to B.L.F.

## References

Axmacher N, Elger CE, Fell J (2008) Ripples in the medial temporal lobe are relevant for human memory consolidation. Brain 131:1806–1817.

Bartoli E, Bosking W, Chen Y, Li Y, Sheth SA, Beauchamp MS, Yoshor D, Foster BL (2019) Functionally Distinct Gamma Range Activity Revealed by Stimulus Tuning in Human Visual Cortex. Curr Biol 29:3345–3358 e3347.

Bragin A, Engel J, Jr., Wilson CL, Fried I, Buzsaki G (1999) High-frequency oscillations in human brain. Hippocampus 9:137–142.

Brainard DH (1997) The Psychophysics Toolbox. Spat Vis 10:433–436.

Brazdil M, Cimbalnik J, Roman R, Shaw DJ, Stead MM, Daniel P, Jurak P, Halamek J (2015) Impact of cognitive stimulation on ripples within human epileptic and non-epileptic hippocampus. BMC Neurosci 16:47.

Buzsaki G (2015) Hippocampal sharp wave-ripple: A cognitive biomarker for episodic memory and planning. Hippocampus 25:1073–1188.

Chen Z, Wilson MA (2017) Deciphering Neural Codes of Memory during Sleep. Trends Neurosci 40:260–275.

Cox RW (1996) AFNI: software for analysis and visualization of functional magnetic resonance neuroimages. Comput Biomed Res 29:162–173.

Dale AM, Fischl B, Sereno MI (1999) Cortical surface-based analysis. I. Segmentation and surface reconstruction. Neuroimage 9:179–194.

Diekelmann S, Born J (2010) The memory function of sleep. Nat Rev Neurosci 11:114–126.

Dudai Y, Karni A, Born J (2015) The Consolidation and Transformation of Memory. Neuron 88:20–32.

Groppe DM, Bickel S, Dykstra AR, Wang X, Megevand P, Mercier MR, Lado FA, Mehta AD, Honey CJ (2017) iELVis: An open source MATLAB toolbox for localizing and visualizing human intracranial electrode data. J Neurosci Methods 281:40–48.

Helfrich RF, Lendner JD, Mander BA, Guillen H, Paff M, Mnatsakanyan L, Vadera S, Walker MP, Lin JJ, Knight RT (2019) Bidirectional prefrontal-hippocampal dynamics organize information transfer during sleep in humans. Nature communications 10:3572.

Iber C, American Academy of Sleep Medicine. (2007) The AASM manual for the scoring of sleep and associated events: rules, terminology and technical specifications. Westchester, IL: American Academy of Sleep Medicine.

Jacobs J, Staba R, Asano E, Otsubo H, Wu JY, Zijlmans M, Mohamed I, Kahane P, Dubeau F, Navarro V, Gotman J (2012) High-frequency oscillations (HFOs) in clinical epilepsy. Prog Neurobiol 98:302–315.

Jiang X, Gonzalez-Martinez J, Halgren E (2019) Coordination of Human Hippocampal Sharpwave Ripples during NREM Sleep with Cortical Theta Bursts, Spindles, Downstates, and Upstates. J Neurosci 39:8744–8761.

Jiang X, Gonzalez-Martinez J, Cash SS, Chauvel P, Gale J, Halgren E (2020) Improved identification and differentiation from epileptiform activity of human hippocampal sharp wave ripples during NREM sleep. Hippocampus 30:610–622.

Joo HR, Frank LM (2018) The hippocampal sharp wave-ripple in memory retrieval for immediate use and consolidation. Nat Rev Neurosci 19:744–757.

Kay K, Frank LM (2019) Three brain states in the hippocampus and cortex. Hippocampus 29:184–238.

Kiani R, Esteky H, Mirpour K, Tanaka K (2007) Object category structure in response patterns of neuronal population in monkey inferior temporal cortex. J Neurophysiol 97:4296–4309.

Klinzing JG, Niethard N, Born J (2019) Mechanisms of systems memory consolidation during sleep. Nat Neurosci 22:1598–1610.

Kumaran D, Hassabis D, McClelland JL (2016) What Learning Systems do Intelligent Agents Need? Complementary Learning Systems Theory Updated. Trends Cogn Sci 20:512–534.

Leonard TK, Hoffman KL (2017) Sharp-Wave Ripples in Primates Are Enhanced near Remembered Visual Objects. Curr Biol 27:257–262.

Leonard TK, Mikkila JM, Eskandar EN, Gerrard JL, Kaping D, Patel SR, Womelsdorf T, Hoffman KL (2015) Sharp Wave Ripples during Visual Exploration in the Primate Hippocampus. J Neurosci 35:14771–14782.

Logothetis NK, Eschenko O, Murayama Y, Augath M, Steudel T, Evrard HC, Besserve M, Oeltermann A (2012) Hippocampal-cortical interaction during periods of subcortical silence. Nature 491:547–553.

McGinley MJ, David SV, McCormick DA (2015a) Cortical Membrane Potential Signature of Optimal States for Sensory Signal Detection. Neuron 87:179–192.

McGinley MJ, Vinck M, Reimer J, Batista-Brito R, Zagha E, Cadwell CR, Tolias AS, Cardin JA, McCormick DA (2015b) Waking State: Rapid Variations Modulate Neural and Behavioral Responses. Neuron 87:1143–1161.

Ngo HV, Fell J, Staresina B (2020) Sleep spindles mediate hippocampal-neocortical coupling during long-duration ripples. Elife 9.

Norman Y, Yeagle EM, Khuvis S, Harel M, Mehta AD, Malach R (2019) Hippocampal sharpwave ripples linked to visual episodic recollection in humans. Science 365.

R Development Core Team (2010) R: A language and environment for statistical computing. In. Vienna, Austria: R foundation for Statistical Computing.

Sirota A, Csicsvari J, Buhl D, Buzsaki G (2003) Communication between neocortex and hippocampus during sleep in rodents. Proc Natl Acad Sci U S A 100:2065–2069.

Staresina BP, Bergmann TO, Bonnefond M, van der Meij R, Jensen O, Deuker L, Elger CE, Axmacher N, Fell J (2015) Hierarchical nesting of slow oscillations, spindles and ripples in the human hippocampus during sleep. Nat Neurosci 18:1679–1686.

Stigliani A, Weiner KS, Grill-Spector K (2015) Temporal Processing Capacity in High-Level Visual Cortex Is Domain Specific. J Neurosci 35:12412–12424.

Vaz AP, Inati SK, Brunel N, Zaghloul KA (2019) Coupled ripple oscillations between the medial temporal lobe and neocortex retrieve human memory. Science 363:975–978.

